# Epigenetic Drift of H3K27me3 in Aging Links Glycolysis to Healthy Longevity

**DOI:** 10.1101/247726

**Authors:** Zaijun Ma, Hui Wang, Yuping Cai, Han Wang, Kongyan Niu, Xiaofen Wu, Huanhuan Ma, Yun Yang, Wenhua Tong, Feng Liu, Zhandong Liu, Yaoyang Zhang, Rui Liu, Zheng-Jiang Zhu, Nan Liu

## Abstract

Epigenetic alteration has been implicated in aging. However, the mechanism by which epigenetic change impacts aging is unclear. H3K27me3, a highly conserved histone modification signifying transcriptional repression, is marked and maintained by Polycomb Repressive Complexes (PRCs). Here, we explore the mechanism by which age-modulated increase of H3K27me3 impacts adult lifespan. Using *Drosophila,* we reveal that aging leads to loss of fidelity in epigenetic marking and drift of H3K27me3 and consequential reduction in the expression of glycolytic genes with negative effects on energy production and redox state. Moreover, we show that a reduction of H3K27me3 by PRCs-deficiency promotes glycolysis and healthy lifespan. While perturbing glycolysis by gene mutation diminishes the pro-lifespan benefits mediated by PRCs-deficiency, transgenic increase of glycolytic genes in wild-type animals extends longevity. Together, we propose that epigenetic drift of H3K27me3 defines a new aging mechanism and that stimulation of glycolysis promotes metabolic health and longevity.

## INTRODUCTION

Aging is characterized by the progressive decline in cellular and organismal functions that lead to reduction of fitness and increased risks to diseases and death. Epigenetic alterations in histone modification represent a prominent hallmark of aging (Sen et al., 2016). Chromatin and epigenetic complexes that modify histones regulate the accessibility of DNA to transcriptional machinery, thereby permitting direct control of gene expression (Kouzarides, 2007). Changes in histone markings during aging have been reported (Liu et al., 2013; Pu et al., 2015; Wood et al., 2010); however, the biological significance of such changes on adult lifespan has not been elucidated. Polycomb repressive complexes (PRCs), including PRC1 (Shao et al., 1999) and PRC2 (Kuzmichev et al., 2002), are important histone-modifying enzymes (Czermin et al., 2002; Muller et al., 2002; Saurin et al., 2001) that are evolutionally conserved from the fruit fly *Drosophila* to mammals (Whitcomb et al., 2007). Tri-methylated histone H3 at lysine 27 (H3K27me3) denotes transcriptional silencing, which is produced by PRC2 (Czermin et al., 2002; Muller et al., 2002) and functionally maintained by PRC1 (Cao et al., 2002). H3K27me3 levels are known to increase with age (Liu et al., 2013; Sun et al., 2014); however, the manner by which H3K27me3 epigenome changes over the course of lifespan and its consequent impact on the progression of aging remain fundamental questions to be addressed.

Metabolic homeostasis is intimately connected with aging and lifespan regulation (Lopez-Otin et al., 2016). Glucose is the dominant provider of energy for cells. Glycolysis, the first step in the breakdown of glucose to extract energy in the form of ATP for cellular metabolism, is present in all cellular organisms and critical for life. The anti-aging effect of caloric restriction correlates with a metabolic feature of limiting glucose metabolism including a reduction of glycolysis (Feuers et al., 1989). On the other hand, glycolysis is essentially involved in a wide-range of biological processes (Chang et al., 2013; Okamoto et al., 2001; Volkenhoff et al., 2015). It is unclear, however, how the beneficial outcome of restricting glucose metabolism and glycolysis as a result of caloric restriction could be reconciled with the essential role of glycolysis during normal adult lifespan. Strikingly, decreased brain glucose metabolism is present prior to the onset of clinic symptoms in individuals at risk of Alzheimer's disease (Cunnane et al., 2011). Since aging is the most significant factor for human neurodegenerative diseases, it is of paramount importance to better understand how glycolysis is modulated with age and whether modulation of glycolytic process could impact adult lifespan.

Mounting evidence associates aging with epigenetic, transcriptional, and metabolic
alterations (Lopez-Otin et al., 2013), yet how these processes are integrated into the regulation of adult physiology is only beginning to be revealed. Though epigenetic effects could potentially elicit broad regulatory mechanisms, it remains to be determined whether epigenetic modulations are simply associated with aging or directly contribute to transcriptional and metabolic modulations that profoundly impact adult lifespan. Here using *Drosophila,* we discover that adult-onset fidelity loss results in epigenetic drift of H3K27me3, which drives the progression of aging by impinging on the transcription and metabolism of the glycolytic pathway. Moreover, we reveal that a reduction of H3K27me3 by PRC2-deficiency promotes glycolysis and healthy lifespan. Our study proposes that stimulation of glycolysis defines a new paradigm for metabolic health and longevity.

## RESULTS

### Adult-onset Fidelity Loss Results in Epigenetic Drift of H3K27me3

H3K27me3 is a signature histone modification, but its dynamics at the epigenome level during aging has not been studied. To characterize H3K27me3 during aging, we examined the level of H3K27me3 by western blot and revealed an age-associated increase in wild-type (WT) flies (Figure S1A). To quantitatively profile H3K27me3, we adapted the chromatin immunoprecipitation (ChIP) followed by high-throughput DNA sequencing with reference exogenous genome method (ChIP-Rx) (Bonhoure et al., 2014; Orlando et al., 2014) in fly tissues (Figure S1B). We generated H3K27me3 epigenome profiles in adult flies of 3d, 15d, and 30d. In addition, we also sampled embryo, larvae, and pupae to examine H3K27me3 epigenome profiles during development. Surprisingly, despite the fact that H3K27me3 levels increased with age, H3K27me3 peak profiles in the aging adult remained largely unchanged (Figure 1A). In a sharp contrast, H3K27me3 signals during developmental stages showed dramatic gain or loss (Figure 1B). This evidence highlights a distinct, yet unknown mechanism of H3K27me3 modification that occurs during aging.

**Figure 1.**
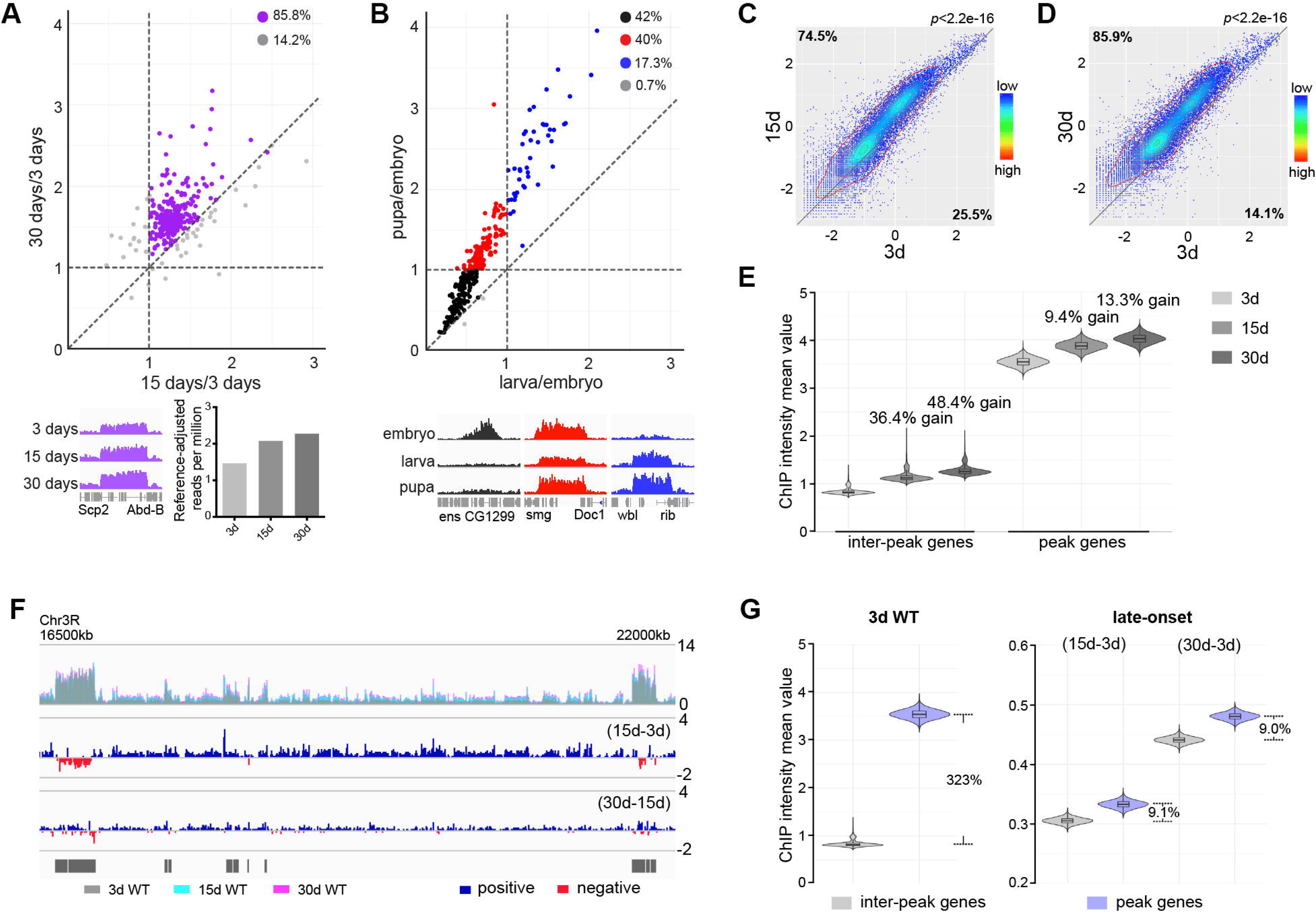
Adult-onset fidelity loss results in epigenetic drift of H3K27me3. (A) H3K27me3 peak profiles are comparable between young and aged animals. A scatter plot (top panel) of 331 peaks and a genome browser mini-view (bottom, left panel) illustrated that H3K27me3 had comparable peak profiles between young and aged animals. Reference-adjusted read counts (bottom, right panel) using ChIP-Rx datasets of 3d, 15d, and 30d exhibited a progressive increase of H3K27me3 signals with age. ChIP-seq was from muscle tissues of 3d, 15d, and 30d old male flies. Genotype: 5905. (B) H3K27me3 modification undergoes dramatic gain or loss during development. A scatter plot (top panel) of 300 peaks showed dynamic changes of H3K27me3 signals during embryo, larvae, and pupae. As illustrated by the genome browser mini-view for select genes (bottom panel), H3K27me3 modification was either embryo-specific (black) or selectively decreased in larvae (red), or progressively increased from embryo, larvae to pupae (blue). ChIP-seq was from whole embryo, larvae and pupae. Genotype as in (A). (C) and (D) H3K27me3 modification increases with age. Scatter plot showed H3K27me3 signals of 17511 genes in 3d as compared to 15d (C) and 30d (D). (wilcoxon signed-rank test, p<2.2e-16). X- and Y-axis represented signal intensity transformed by Log2. Contour lines indicated that H3K27me3 signals were higher in aged flies compared to 3d old flies. ChIP-seq and genotype as in (A). (E) Inter-peak genes receive relatively more H3K27me3 modification during aging. Violin plots for inter-peak genes (left) and peak genes (right) at 3d, 15d, and 30d of age. We computed bootstrapped confidence intervals for the mean ChIP intensity (10,000 draws with replacement of n=500). Net gain of H3K27me3 signals during aging was calculated by a subtraction of the mean signal intensity between aged and 3d. Bootstrapped 95% confidence intervals: 3d inter-peak genes: [0.758, 1.017]; 15d inter-peak genes: [1.023, 1.401]; 30d inter-peak genes: [1.149, 1.562]; 3d peak genes: [3.349, 3.739]; 15d peak genes: [3.681, 4.075]; 30d peak genes: [3.817, 4.230]. ChIP-seq and genotype as in (A). (F) H3K27me3 modification during aging has reduced selectivity. Genome browser view of a 5.5Mb region in the chromosome 3R was shown. H3K27me3 occupancy overlaid between 3d (grey), 15d (cyan), and 30d (pink) (top panel). H3K27me3 modification during aging was shown by deducting 3d signals from those at 15d (middle panel) and by deducting 15d signals from those at 30d of age (bottom panel). Black boxes and lines represented peak regions, corresponding to their chromosomal locations. ChIP-seq and genotype as in (A). (G) Violin plots indicate that signals are highly preferential at the peak regions at 3d (left), but for signals gained during aging, selectivity is dramatically reduced (right). Bootstrapped 95% confidence intervals: 3d inter-peak genes: [0.758, 1.017]; 3d peak genes: [3.349, 3.739]; late-onset (15d-3d) inter-peak genes: [0.295, 0.315]; (15d-3d) peak genes: [0.320, 0.345]; (30d-3d) inter-peak genes: [0.430, 0.452]; (30d-3d) peak genes: [0.468, 0.493]. ChIP-seq and genotype as in (A).

To define the landscape of aging epigenome, we divided distribution of H3K27me3 into peak regions (IP/input ≥2 and signal spanning≥3kb) and inter-peak regions. Analysis in 3d WT revealed 222 peak regions reproducibly identified from a total of four replicates, which covered approximately 9.73 Mbp of the genome (Figure S1C; Table S1). Analysis of global occupancy of H3K27me3 between young and aged WT exhibited similar patterns (pair-wise Pearson correlation coefficients ≥0.70) (Figure S1D), implying that the genome-wide distributions of H3K27me3, particularly the peak regions, are well maintained in aged WT.

We next analyzed signal intensity of individual genes using reference-adjusted read counts. Scatter plots demonstrated that H3K27me3 signals in WT clearly had an adult-onset, progressive increase (Figures 1C and 1D). To further characterize H3K27me3 dynamics, we sorted individual genes into peak genes (1141 genes) according to the peak regions and genes therein covered (see Table S1 for peak regions and included genes) or inter-peak genes (16370 genes). In aging WT flies, despite a genome-wide increase, we noted that inter-peak genes gained relatively more signals than those of peak genes (Figure 1E). To quantify the H3K27me3 marks gained specifically during aging, we subtracted the signals of 3d from those of older age points. Genome browser views illustrated that subtracted signals propagated along the chromosome, showing no direct correlation with pre-existing peaks (Figure 1F). We then extended this analysis to the entire epigenome. Modification in flies that were 3d old was predominantly biased at the peaks, with approximately 323% more signals as compared to the inter-peaks (Figure 1G). In contrast, the signals acquired during aging were only about 9% more at the peaks than those at the inter-peaks (Figure 1G), suggesting that there was a dramatic shift in the pattern of H3K27me3 modification in aging.

Altogether, analysis of H3K27me3 aging dynamics implicates a strong selection on peak regions compared to inter-peak regions when animals are young; however, the fidelity of this regulation becomes less stringent during aging, and consequently, leading to the epigenetic drifting of H3K27me3 modification into the much broader inter-peak regions as observed in aged animals.

### A CRISPR/Cas9 Deficiency Screen Identifies the Role of PRCs in Aging

Given above study indicated a new feature of H3K27me3 dynamics at the epigenome level during adult lifespan, we further explored the roles of epigenetic pathways in the aging process via an unbiased deficiency-based genetic screen. Using CRISPR/Cas9 mutagenesis (Ren et al., 2013b), we systematically generated site-specific deletion mutants of 24 key regulators involved in distinct epigenetic and chromatin modifications in the same homogenous genetic background (Table S2). Since the effects of epigenetic genes are usually dose-dependent, we analyzed adult survival using heterozygous mutants. Lifespan analysis demonstrated that whereas majority of mutants showed mild or no effect, animals deficient in PRCs, including *esc, E(z), Pcl, Su(z)12* of PRC2 and *Psc* and *Su(z)2* of PRC1, lived substantially longer (Figure 2A; Table S2). We further examined pairwise combinations of PRC2 double mutants in trans-heterozygosity. Remarkably, double mutants demonstrated more striking effects on H3K27me3-reduction and life-extension (Figure 2B; Figure S2A). Identification of multiple components of PRC2 and PRC1 in the screen is interesting, suggesting the involvement of the epigenetic changes regulated by these two complexes in animal lifespan.

**Figure 2.**
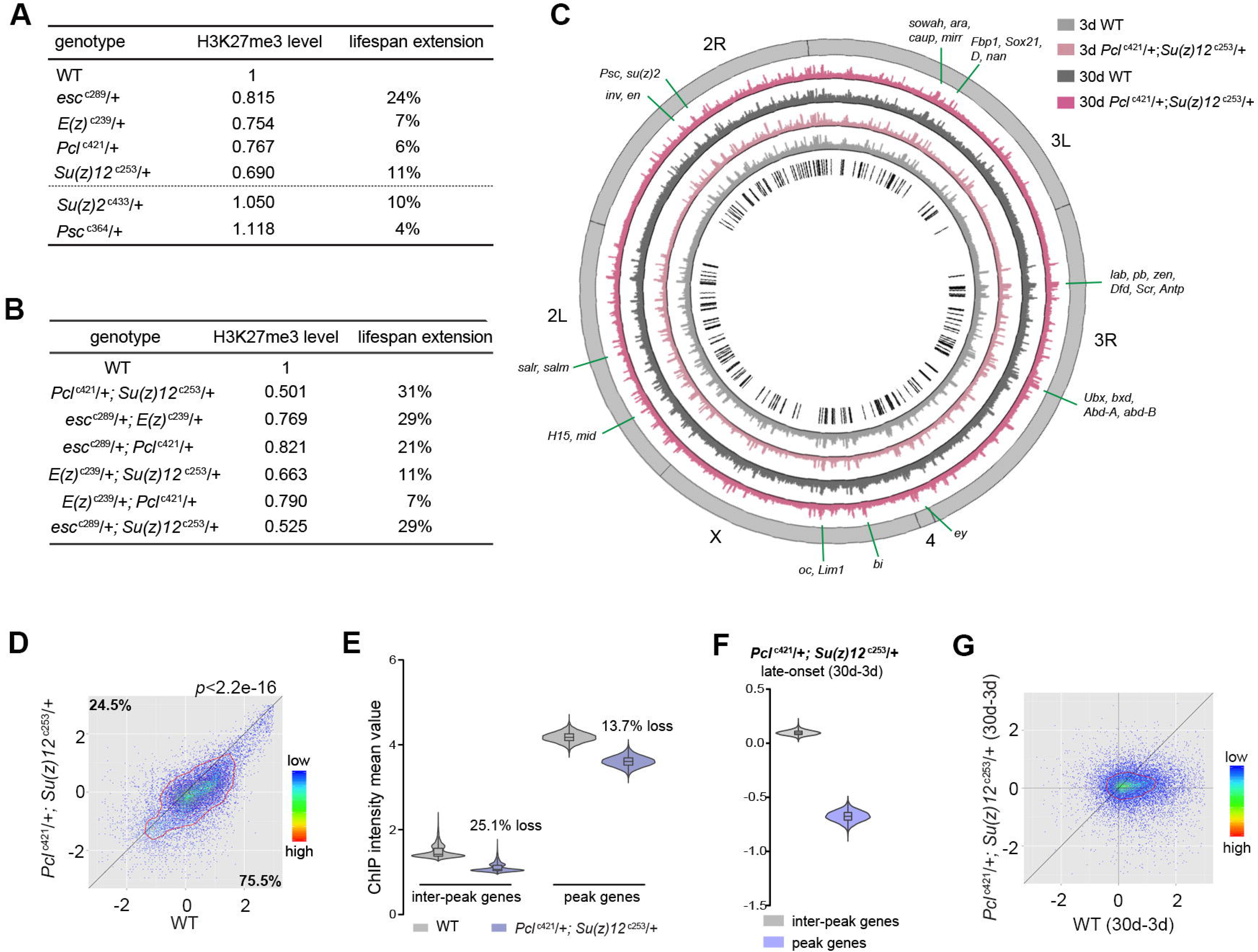
Long-lived PRC2 mutants diminish the epigenetic drift of H3K27me3 during aging. (A) PRCs-deficient animals have extended lifespan. PRC2 heterozygous mutants of indicated genotype reduce H3K27me3 levels and extend lifespan (top of the dashed line); PRC1 heterozygous mutants of indicated genotype extend lifespan without changing H3K27me3 levels (bottom of the dashed line). To name new mutant, a superscript amended to the gene contained a letter c denoting CRISPR/Cas9 method followed by the size of genomic deletion. All mutants have been backcrossed with WT for five times to ensure a uniform genetic background. See also Table S2. Western blot was from head tissues of 3d old male flies. (for H3K27me3 quantification: mean±SD of 3 biological repeats; student *t*-test; for lifespan assay: 25°C; n≥200 per genotype for curve; log-rank test). (B) Pair-wise combination of PRC2 trans-heterozygous double mutants of indicated genotype results in stronger effects in H3K27me3-reduction and life-extension. See also Figure S2A. (C) Circos plot of the H3K27me3 epigenome illustrates peak profiles that are highly preserved with age and in PRC2 mutants. Black boxes and lines (innermost circle) represented common peak regions, corresponding to their chromosomal locations. Chromosome ideogram was in grey (outermost ring). PRC2 target genes previously found in cells and during development were shown next to their epigenomic loci. ChIP-seq was from muscle tissues of 3d and 30d old male flies. Genotypes: WT: 5905 and *Pcl*^c421^/+; *Su(z)12*^c253^/+. (D) H3K27me3 modification decreases in PRC2 mutants. A scatter plot showed H3K27me3 signals of 17511 genes in *Pcl*^c421^; *Su(z)12*^c253^ as compared to WT. (wilcoxon signed-rank test, p<2.2e-16). X- and Y-axis represented signal intensity transformed by Log2. Contour lines indicated that H3K27me3 signals were higher in WT compared to mutants. ChIP-seq was from muscle tissues of 3d old male flies. Genotypes: WT: 5905 and *Pcl*^c421^/+; *Su(z)12*^c253^/+. (E) Inter-peak genes receive relatively less H3K27me3 modification in PRC2 mutants. Reduction of signals was calculated by a subtraction of the mean signal intensity between WT and PRC2 mutants. Bootstrapped 95% confidence intervals: 3d inter-peak genes: [1.317, 1.861]; 3d *Pcl*^c421^; *Su(z)12*^c253^ inter-peak genes: [0.944, 1.361]; 3d peak genes: [3.946, 4.414]; 3d *Pcl*^421^; *Su(z)12*^253^ peak genes: [3.364, 3.861]. ChIP-seq and genotypes as in (D). (F) Modification of H3K27me3 during aging has reduced selectivity in PRC2 mutants. Bootstrapped 95% confidence intervals: *Pcl^421^*; *Su(z)12^253^* late-onset inter-peak genes: [0.053, 0.147]; *Pcl^421^; Su(z)12^253^* late-onset peak genes: [-0.778, −0.578]. ChIP-seq was from muscle tissues of 3d and 30d old male flies. Genotypes: *Pcl*^c421^/+; *Su(z)12*^c253^/+. (G) Age-associated drifting of H3K27me3 is dampened in PRC2 mutants. Scatter plot for 17511 genes between WT and mutants. X- and Y-axis represented signal intensity transformed by Log2. Contour lines indicated that the majority of the gene signals displayed higher levels—thus more rapid changes—in aged WT compared to mutants. ChIP-seq was from muscle tissues of 3d and 30d old male flies. ChIP-seq and genotypes as in (D).

### Long-lived PRC2 Mutants Diminish the Epigenetic Drift of H3K27me3 During Aging

We then investigated whether the age-delaying effect of PRCs-deficiency might be due to its ability to modulate H3K27me3 dynamics with age. To address this, we examined the modification of H3K27me3 using young (3d) and aged (30d) mutants with their age-matched WT controls. For this and subsequent experiments, we studied the PRC2 deficiency using *Pcl*^c421^ *Su(z)12*^c253^ trans-heterozygote double mutants, which gave rise to the strongest effect in H3K27me3-reduction and life-extension. Of note, western blot analysis showed that only H3K27me2/3 were selectively reduced in PRC2 mutants, while other histone markings remained unchanged (Figure S2B). Interestingly, global occupancy of H3K27me3 between WT and age-matched PRC2 mutants was highly preserved, as demonstrated by the similar patterns of peak profiles and genes known to be H3K27me3-modified (pair-wise Pearson correlation coefficients ≥0.77) (Figure 2C; Figures S2C and S2D). H3K27me3 signals in WT were uniformly higher than those in mutants (Figure 2D; see Figure S2E for *esc*^c289^ *E(z)*^c239^). In PRC2 mutants at 3d where H3K27me3 bulk levels were substantially decreased, interestingly, the signals became more likely to coalesce into the peak regions (Figure 2E; see Figure S2F for *esc*^c289^ *E(z)*^c239^). Given limited H3K27me3 repertoire in the mutants, this shift might be critical for the establishment of the peaks. In aging PRC2 mutants, adult-onset modification was less biased or even decreased at the peaks (Figure 2F; see Figure S2G for *esc*^c289^*E(z)*^c239^). More strikingly, age-associated shift in H3K27me3 modification that occurred in WT was now dampened in mutants (Figure 2G; see Figure S2H for *esc*^c289^*E(Z)*^c239^). Combined, our study implicates that the life-extension effect of PRC2-mutation likely arises from its ability to mitigate the drifting of H3K27me3 during aging.

### Transcriptomics Links H3K27me3 Dynamics to the Regulation of Glycolytic Genes

Since biological activities of histone modification often link chromatin accessibility to gene expression (Kouzarides, 2007), we speculated that alterations in the levels of H3K27me3, a repressive epigenetic mark in transcription, could impact the transcriptome. We asked whether transcriptional change of particular genes might account for PRC2-dependent life-extension. Using muscle tissues, we generated RNA-seq datasets for polyA-selected mRNAs. Analysis of individual PRC2 mutants resulted in several hundreds of upregulated genes, most of which were those at the inter-peak regions (Figure S3A). On the other hand, analysis of 63 genes with known effects on aging, including genes in insulin/IGF-1, mTOR pathways, etc., revealed no consistent changes in the expression in PRC2 mutants (Table S3), thus excluding them from being the major contributors in mediating PRC2-dependent longevity. Interestingly, comparative transcriptomics converged on a common set of genes (Figure 3A; Table S3). Gene ontology (GO) analysis revealed that the “glycolytic process” and closely related pathways were highlighted for genes upregulated (Figure 3A), while the “oxidation-reduction process” was enriched for genes downregulated (Figure 3A). Using weighted gene co-expression network analysis (WGCNA) (Langfelder and Horvath, 2008), we determined the significance of highly correlated changes in the expression of glycolytic genes (Figure 3B). We then extended RNA-seq analysis to the heads, and further narrowed down to two glycolytic genes, *Tpi* and *Pgi*, whose expressions were upregulated in both tissue types across individual long-lived PRC2 mutants (Figure S3B; Table S4). As noted, glucose is metabolized through sequential reactions of glycolysis and the citric acid cycle. However, we found no consistent changes in the expression of genes mediating pentose phosphate pathway (PPP), citric acid cycle, and oxidative phosphorylation (Figure S3B; Table S4), suggesting modulatory effects specifically on the glycolytic genes.

**Figure 3.**
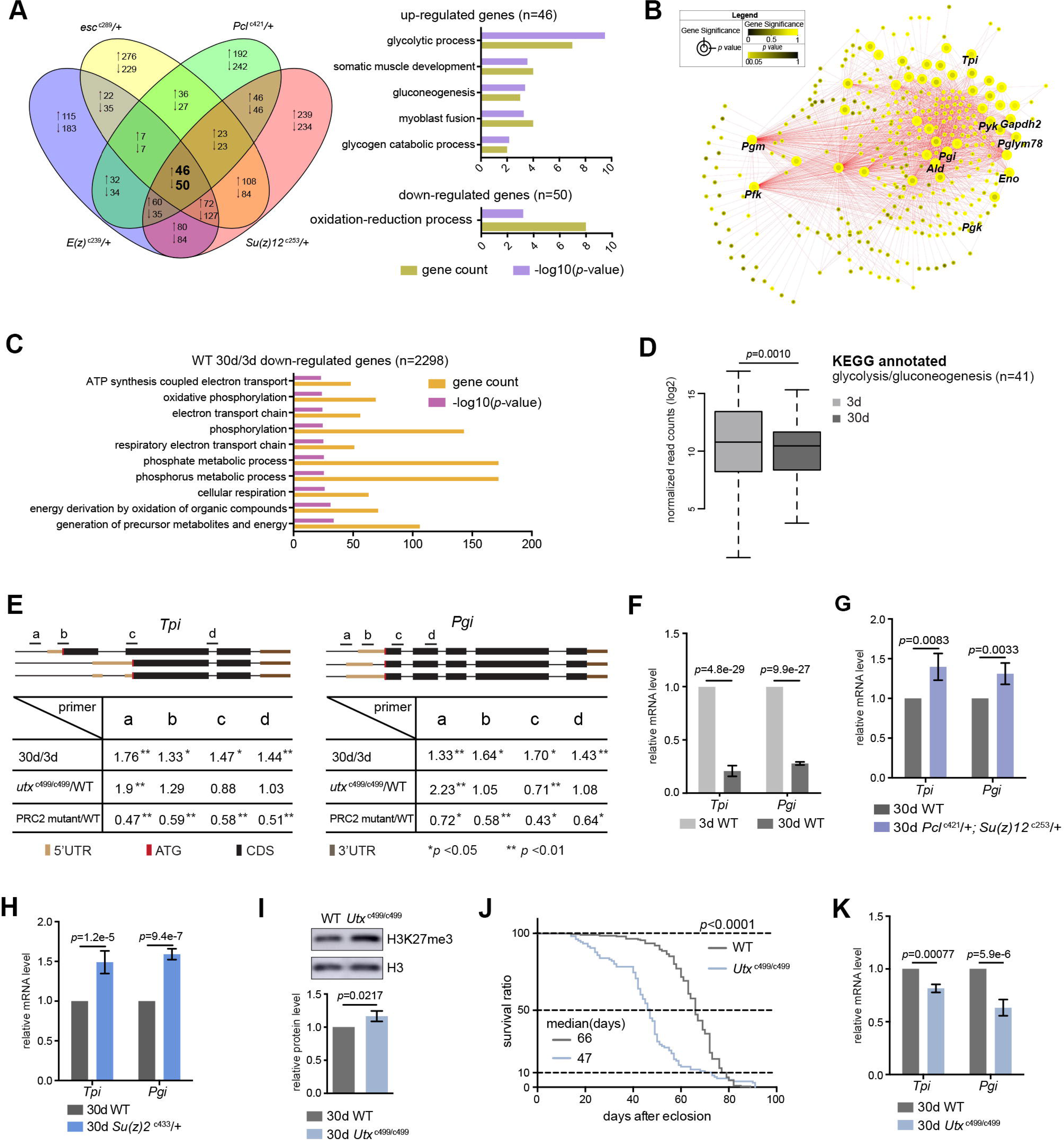
Transcriptomics links H3K27me3 dynamics to the regulation of glycolytic genes. (A) Venn diagram shows genes commonly changed in PRC2 single mutants with indicated genotype (left panel). GO analysis (right panel) shows glycolysis being the biological processes significantly enriched for genes upregulated in PRC2 long-lived mutants, while oxidation-reduction process is the only pathway enriched for genes down-regulated. (B) WGCNA network analysis reveals a co-regulated change of glycolytic genes. Glycolytic genes were highlighted. (C) GO analysis ranks biological processes related to energy metabolism being significantly decreased with age. (2298 genes were decreased with age by using a cutoff of p<0.05). The top 10 most significantly affected biological processes were shown. RNA-seq was from muscle tissues of 3d and 30d old male flies. Genotype: 5905. (D) Genes of glycolysis/gluconeogenesis as annotated in the Kyoto Encyclopedia of Genes and Genomes (KEGG) pathway database show age-associated transcriptional decrease. Y-axis represented normalized read counts transformed by Log2. (41 genes used; see Table S4; wilcoxon signed rank test). RNA-seq and genotype as in (C). (E) ChIP-qPCR validation. Genomic structures were shown according to the Flybase annotation (https://www.flybase.org). Along the gene of indicated genotype, a, b, c, and d were underlined, denoting the sites for PCR amplification. Color codes represented ATG (the translation start site), CDS (coding sequence), 5’UTR, and 3’UTR. Different primer sets confirmed that H3K27me3 modification was increased in aged WT and further increased in *utx* null mutants. H3K27me3 was reduced in PRC2 mutants as compared to WT, with the ratio between mutants and WT smaller than 1 (mean±SD of 3 biological repeats; student *t*-test *p<0.05, **p <0.01; also see Figure S3C). ChIP-qPCR was from muscle tissues of 3d and 30d old male flies. Genotypes: WT: 5905. PRC2 mutant: *Pcl*^421^/+; *Su(z)12*^c253^/+. *utx*^c499/c499^. (F-H) qRT-PCR analysis confirms that *Tpi* and *Pgi* of glycolytic genes have a decrease with age (F) but become increased in both PRC2 (G) and PRC1 mutants (H). (mean±SD of 3 biological repeats; student t-test). qRT-PCR was from muscle tissues of 3d and 30d old male flies. Genotypes: WT: 5905. *Pcl*^c421^/+; *Su(z)12*^c253^/+. *Su(z)2*^c433^/+. (I) H3K27me3 increases in *utx* deficiency. H3K27me3 western blot (top) and quantification (bottom). (for H3K27me3 quantification: mean±SD of 3 biological repeats; student t-test). Western blot was from muscle tissues of 30d old male flies. Genotypes: WT: 5905. *utx*^c499/c499^. (J) *utx* null animals are short-lived. (for lifespan assay: 25°C; n≥200 per genotype for curve; log-rank test). Genotypes as in (I). (K) qRT-PCR analysis indicates a further decrease of glycolytic genes in *utx* deficient animals. (mean±SD of 3 biological repeats; student *t*-test). qRT-PCR was from muscle tissues of 30d old male flies. Genotypes as in (I).

Moreover, RNA-seq analyses of WT animals revealed that, though aging might exert a much broader effect on the changes of the transcriptome, genes downregulated with age were mainly enriched in carbohydrate metabolism and energy production (Figure 3C). In particular, we noted a transcriptional decline of genes related to glycolysis/gluconeogenesis in old animals (Figure 3D). To verify these results, we subsequently performed single-gene analysis on *Tpi* and *Pgi* with respect to their H3K27me3 states and mRNA levels using ChIP-qPCR (Figure 3E; Figure S3C) and qRT-PCR (Figures 3F and 3G), respectively. In addition to PRC2 mutants, we attested the transcriptional increase of these two glycolytic genes in *Su(Z)2*^c433^, a long-lived PRC1 mutant (Figure 3H).

Given the fact that H3K27me3 naturally increased with age, we investigated whether further increase of H3K27me3 could impact lifespan and gene expression. To address this, we examined flies deficient in histone demethylation. The *Drosophila Utx* gene has been previously implicated in catalyzing the removal of methyl groups from H3K27me3 (Smith et al., 2008); a null mutation, *utx*^c499^ (Table S2), displayed a further increase of H3K27me3 compared to age-matched WT (Figures 3E and 3I; Figure S3C). Flies lacking *utx* were viable, but detailed characterization indicated a significantly shortened lifespan, to about 70% that of WT animals (Figure 3J). Concomitant with an increase in H3K27me3 modification, expression of both *Tpi* and *Pgi* was further decreased (Figure 3K). Finally, control studies using short-lived *miR-34* mutants (Liu et al., 2012) and a long-lived *piwi* deficiency that was identified from our unbiased genetic screen (Table S2) demonstrated that transcriptional changes of the glycolytic genes had no simple correlation with lifespan alterations (Figures S3D and S3E).

Altogether, these data reveal that H3K27me3 dynamics with natural aging causes a downregulation of glycolytic genes, and this reduction is enhanced in *utx* mutants upon further increase of H3K27me3. Long-lived PRC2 mutants confer a specific effect by reversing the decline in the expression of glycolytic genes that are otherwise significantly decreased with age.

### Untargeted Metabolomics Shows Enhanced Glycolysis in Long-lived PRC2 Mutants

To further investigate the mechanism by which epigenetic changes regulate aging, we conducted another unbiased study by profiling cellular metabolites using LC-MS based untargeted metabolomics (Patti et al., 2012). In total, 151 metabolites were identified. This experiment demonstrated changes in the levels of metabolites during aging of WT flies, and between WT and mutants (Figures 4A and 4C; Figure S4A). Pathway analysis revealed that glycolysis among additional metabolic processes was significantly enriched (Figures 4B and 4D). In particular, lactate, a specific indicator for anaerobic glycolysis (Schurr and Payne, 2007), was significantly decreased during normal aging in WT animals but became elevated in PRC2 mutants (Figure 4E). Of note, metabolites related to citric acid cycle remained unchanged between WT and mutants (Figure S4B). Combined, this evidence reveals a metabolic profile reflective of modulated glycolysis, being decreased during normal aging but increased in long-lived PRC2 mutants. Thus, this unbiased study using metabolomic approach showed modulation in the level of lactate and glycolysis during aging and in PRC2 mutants, in good congruence with the conclusion from the studies of epigenomic and transcriptional alterations (Figures S3B and S3C).

**Figure 4.**
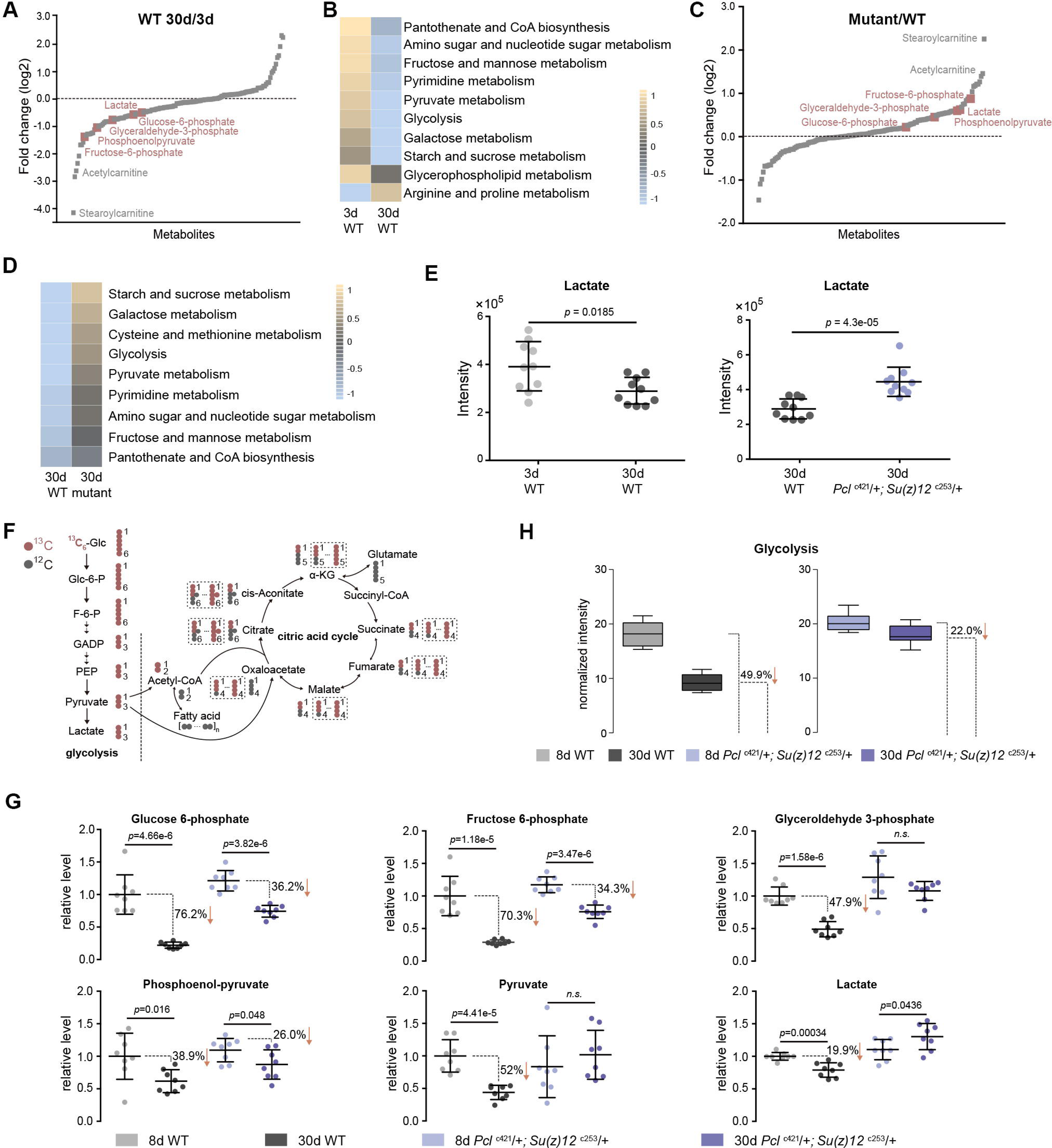
Metabolomics shows the effect of PRC2 mutants in reversing glycolytic decline in aging. (A) Untargeted metabolics identifies that metabolites of the glycolytic pathway (red) are decreased with age. For each metabolite, the median from 10 biological repeats was used to calculate the relative fold change between WT and mutants. Untargeted metabolics was from head tissues of 3d and 30d old male flies. Genotype: 5905. (B) Pathway enrichment analysis reveals glycolysis being the biological processes significantly enriched with age. (*p*<0.01; see STAR method for algorithm). Genotype as in (A). (C) Untargeted metabolics identifies that metabolites of the glycolytic pathway (red) are increased in PRC2 mutants. Untargeted metabolics and analysis as in (A). Genotypes: WT: 5905. Mut: *Pc*^c421^/+; *Su(z)12*^cZ53^/+. (D) Pathway enrichment analysis reveals glycolysis being the biological processes significantly enriched. (p<0.01; see STAR method for algorithm). Genotype as in (C). (E) Lactate is decreased with age (left panel) but increased in PRC2 mutants (right panel). (mean±SD of 10 biological repeats; wilcoxon rank-sum test). Genotypes: WT: 5905. *Pcl*^c421^/+; *Su(z)12*^c253^/+. (F) Schematic representation of the glucose flux via glycolysis (left) and citric acid cycle (right). ^13^C and ^12^C were indicated for specific metabolites. Dashed rectangles encircled isotopologues pertaining to specific metabolites that contained different number of ^13^C. (G) Decline of glycolytic metabolites during aging is partially rescued by PRC2 deficiency. For specific metabolite, the median from 8 biological repeats was used to calculate the ratio between 8d and 30d of age. (mean±SD of 8 biological repeats; wilcoxon rank-sum test). Metabolomics was from head tissues of 8d and 30d old male flies. Genotypes: WT: 5905. *Pcl*^c421^/+; *Su(z)12*^c253^/+. (H) Age-associated decline of glycolysis is diminished by PRC2 deficiency. ^13^C-traced glycolytic metabolites were used to determine the level of glycolytic pathway as a whole. In 30d WT, 50.1% of glycolysis remained as relative to its level at 8d of age; in contrast, in PRC2 mutants at 30d of age, 78% of glycolysis remained. (mean±SD of 8 biological repeat; see STAR method for calculation). Metabolomics and genotypes as in (G).

### Metabolic Flux Shows Roles of PRC2 Mutants in Reversing Glycolytic Decline in Aging

To provide quantitative insights into the changes in glucose metabolism with age and in PRC2 mutants, we devised a metabolic flux assay using ^13^C-labeled glucose as a tracer in adult aging flies. For young (3d) and aged (25d) animals, fly diet was switched from ^12^C-glucose to ^13^C-glucose, such that the glucose metabolism can be traced by measuring the ^13^C-labeled metabolites (Figure 4F; Figure S5A). After 5 days of feeding, ^13^C-labeled glucose accounted for more than 93% of the total glucose pool (Figure S5B), suggesting a near-complete replacement. This experiment demonstrated that while the level of labeled glucose was similar (Figure S5C), specific metabolites (Figure 4G) and the glycolytic pathway as a whole (Figure 4H) exhibited an age-associated decline; strikingly, the extent of decline was partially reversed by PRC2-deficiency. This finding, together with above results obtained by untargeted metabolomics, provides definitive evidence that glycolysis is modulated by aging. Since metabolites of the citric acid cycle are mix lineages from glycolysis, glutamate, and fatty acids, each metabolite may contain multiple ^13^C-labeled isotopologues via glycolysis-derived reactions as well as recurrent cycling (Figure 4F). Thus we analyzed all isotopologues pertaining to each metabolites (Figure S5D) and their summed intensity (Figure S5E). Our data indicated no consistent change of the citric acid cycle with age and in PRC2-deficiency, which was in line with constant levels of metabolites as above measured by untargeted metabolomics (see Figure S4B).

These data combined establish a regulatory event from epigenetic alterations to transcriptional and metabolic modulations impacting the manner by which glucose can be metabolized during aging and in PRC2 mutants. While natural aging leads to a metabolic decline including a reduction of glycolysis, PRC2 deficiency stimulates glycolysis, thereby permitting a proper level of glycolysis to be maintained with age.

### Stimulation of Glycolysis Promotes Energy, Redox State and Adult Fitness

We next characterized the contributions of specific metabolites in the glycolytic pathway. While anaerobic glycolysis yields ATP only, glycolysis at the step of pyruvate formation has a net outcome of both ATP and NADH, a reduced form of nicotinamide adenine dinucleotide (NAD^+^) (see Figure S3B). While the overall glucose levels between WT and mutants had no significant difference (Figure 5A), pyruvate was increased in mutants (Figure 5A). On the other hand, the ratio of NADPH/NADP^+^, an indicator of the PPP, foliate metabolism, and malic enzyme (Fan et al., 2014) was even slightly decreased in mutants (Figure 5B). In parallel, we also examined the glutathione, an important indicator of the cellular redox state (Jones, 2006). Our data showed that ATP, NADH/NAD^+^, and GSH/(GSH+GSSG) ratio were decreased with age but increased in mutants (Figures 5C and 5D). Combined, analysis of specific metabolites demonstrates a metabolic decline associated with normal aging, and that stimulation of glycolysis in PRC2 mutants promotes energy and cellular redox potential.

**Figure 5.**
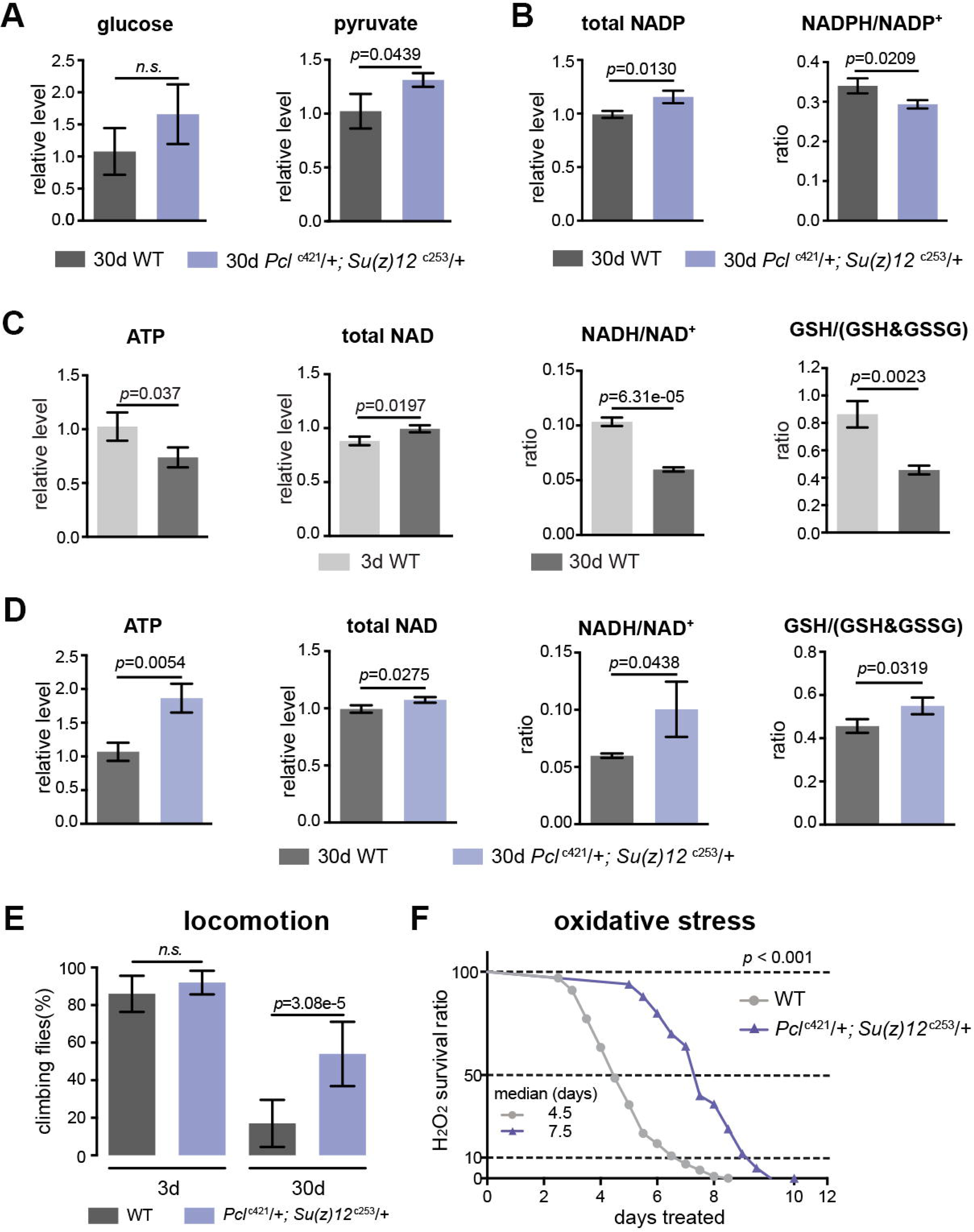
PRC2 mutants couple enhanced glycolysis with improved adult fitness. (A) Glucose content shows a slight increase in PRC2 mutants, but this increase is not statistically significant (left panel). Pyruvate, one key end product of glycolysis, is increased (right panel). (mean±SD of 3 biological repeats with 10 flies for each measurements; student *t*-test; *n.s.:* not significant). Test was from muscle tissues of 30d old male flies. Genotypes: WT: 5905. Mut: *Pcl*^c421^/+; *Su(z)1*2^c253^/+. (B) The ratio of NADPH/NADP^+^, indicator of the PPP, foliate metabolism, and malic enzyme is decreased in mutants. Test, statistics, and genotype as in (A). (C) ATP and cellular redox levels are decreased with age. (mean±SD of 3 biological repeats with 10 flies for each measurements; student *t*-test). Test was from muscle tissues of 3d and 30d old male flies. Genotype: 5905. (D) ATP and cellular redox levels are increased in PRC2 deficient animals. Test, statistics, and genotype as in (A). (E) and (F) Analysis of adult phenotypes reveals that PRC2 mutants have a healthy lifespan. Climbing assay exhibited that, whereas WT and mutants behaved similarly at 3d, with age, mutants had better climbing, reflective of improved mobility (E). Mutants had enhanced resistance to oxidation (F). (for climbing assay: mean±SD of 10 biological repeats with 10 flies for each repeats; student *t*-test; for oxidation tests: 25°C; n=100 per genotype for curve; log-rank test). Genotypes as in (A).

We then asked how this ability of enhanced glycolysis in PRC2 mutants was translated into fitness and stress resistance in aging. To illuminate this, we characterized age-related phenotypes. Analysis of climbing of young (3d) and aged (30d) animals demonstrated that whereas WT and mutants at 3d behaved similarly, PRC2 mutants at 30d of age had better locomotion (Figure 5E). To examine the sensitivity to oxidative stress, we utilized hydrogen peroxide (H_2_O_2_), a source of reactive oxygen species (ROS). We observed substantially enhanced resistance to oxidation in mutants (Figure 5F). Elevated energy and NADH/NAD^+^ ratio in PRC2 mutants might underlie improved motility and resistance to ROS, respectively. These data implicate that PRC2-deficiency couples longevity and metabolism with enhanced ability to handle stress, a property reflecting improved adult fitness.

### Perturbing Glycolysis Diminishes Longevity Traits in PRC2 Mutants

To ask if the activation of glycolysis may directly contribute to healthy lifespan, we determined whether perturbing glycolysis could diminish the lifespan benefits of PRC2 mutants. In PRC2 mutants, *Tpi* and *Pgi* were two genes exhibiting a concerted decrease in H3K27me3 modification (see Figure 3E; Figure S3C) and a corresponding increase in transcription (see Figure 3G; Figure S3B); thus *Tpi* and *Pgi* might be important targets of PRC2. In light of this, we generated a null mutation of the *Tpi* gene by CRISPR/Cas9, *Tpi*^c511^ (Table S2). *Tpi* gene, encoding triosephosphate isomerase, is a central glycolytic enzyme (Bar-Even et al., 2012). Given homozygous *Tpi*^c511^ had a pre-adult lethality, we combined *Tpi*^c511^ heterozygote with *Pcl*^c421^ *Su(z)12*^c253^, and evaluated the adult phenotypes by using the triple mutants. Control studies indicated that the *Tpi*^c511^ heterozygote alone retained normal lifespan (Figure S6A), oxidative stress (Figure S6B), locomotion (Figure S6C), and comparable levels of ATP and NADH/NAD^+^ ratio (Figure S6D). Lowering *Tpi* in PRC2 mutants selectively diminished the extent by which glycolysis could be stimulated as having relatively reduced ATP and NADH/NAD^+^ ratio (Figure 6D). Analysis of lifespan exhibited that *Tpi* deficiency partially mitigated the longevity phenotype in PRC2 mutants (Figure 6A). Traits related to adult fitness, including resistance to ROS and climbing mobility, were also attenuated by *Tpi* deficiency (Figures 6B and 6C). To consolidate this result, we generated a null mutation for *Pgi* by CRISPR/Cas9, *Pgi*^c392^ (Table S2). Consistently, while *Pgi*^c392^ heterozygote alone had minimal effect on aging (Figures S6E-S6H), triple mutants combining *Pgi*^c392^ heterozygote with *Pcl*^c421^ *Su(z)l2*^c253^ diminished the pro-lifespan benefits mediated by PRC2-deficiency (Figures 6E-6H). These data implicate that the life-extension effects of PRC2 mutants require the activation of glycolysis.

**Figure 6.**
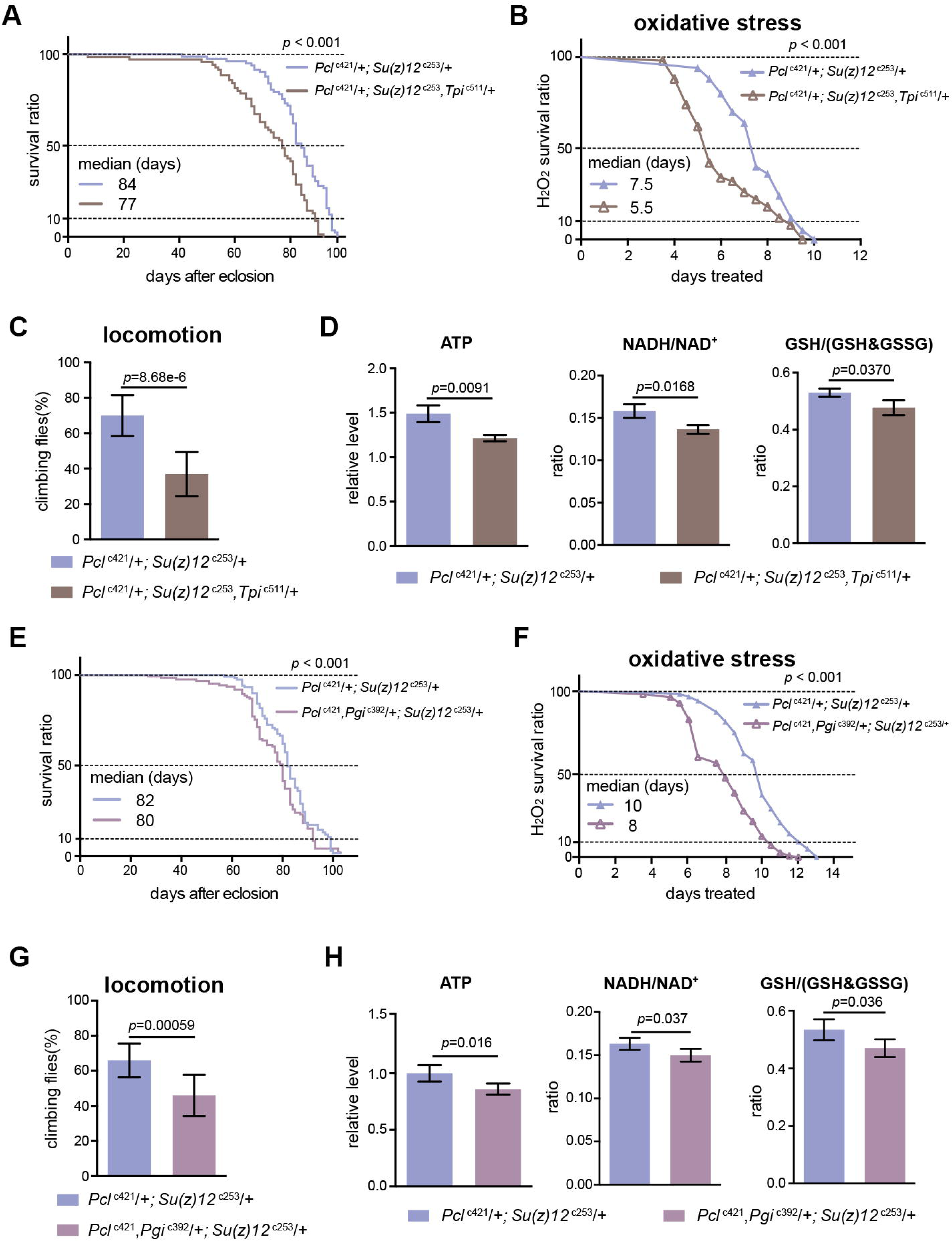
Perturbing glycolysis diminishes longevity benefits in PRC2 mutants. (A) *Tpi* deficiency significantly diminishes the longevity phenotype of PRC2 trans-heterozygous double mutants. (for lifespan assay: 25°C; n>200 per genotype; log-rank test). Genotypes: *Pcl*^c421^/+; *Su(z)12*^c253^/+;. *Pcl*^c421^/+; *Su(z)12*^c253^, *Tpi*^c511^/+. (B) and (C) Oxidation stress test (B) and climbing (C) show that *Tpi* deficiency partially diminishes the lifespan-benefits of PRC2 mutants. (for oxidation assay: 25°C; n=100 per genotype for curve; log-rank test; for climbing assay: mean±SD of 10 biological repeats with 10 flies for each repeat; student *t*-test). Genotypes as in (A). (D) *Tpi* deficiency reduces glycolysis of PRC2 mutants. Analysis of specific metabolites revealed a decrease of ATP (left panel), ratio of NADH/NAD^+^ (middle panel), and ratio of GSH/(GSH+GSSG) (right panel). (mean±SD of 3 biological repeats; student *t*-test). Metabolite analysis was from muscle tissues of 30d old male flies. Genotypes as in (A). (E-H) *Pgi* deficiency diminishes the aging benefits mediated by PRC2 mutants, including lifespan (E), oxidative stress (F), locomotion (G), and glycolysis (H). Genotypes: *Pcl*^c421^/+; *Su(z)12*^c253^/+. *Pcl*^c421^, *Pgl*^c392^/+; *Su(z)12*^c253^/+.

### Transgenic Increase of Glycolytic Genes Promotes Healthy Lifespan

Our data thus far have established the role of glycolysis in PRC2-dependent life-benefits. Furthermore, we investigated whether upregulating glycolytic genes in WT animals could promote glycolysis, which in turn extend lifespan. To address this, we increased the gene dosage of *Tpi* and *Pgi* via genomic transgenes, which expressed these genes under their endogenous regulatory elements (Figures S7A and S7B). Western blot analysis manifested that both Tpi and Pgi proteins had a modest decrease with age, but became upregulated upon PRC2-deficiency, an outcome consistent with the alteration in the level of H3K27me3 (Figures 7A-7D). We then interrogated the effects of *Tpi* and *Pgi* transgenes on WT flies. While single transgenes had modest phenotypes, animals combining both *Tpi* and *Pgi* transgenes significantly stimulated glycolysis, as shown by elevated pyruvate, ATP, and NADH/NAD^+^ ratio compared to age-matched WT (Figure 7E). Accordingly, adult aging phenotypes, including lifespan (Figure 7F), locomotion (Figure 7G), and resistance to oxidative stress (Figure 7H), were substantially improved.

**Figure 7.**
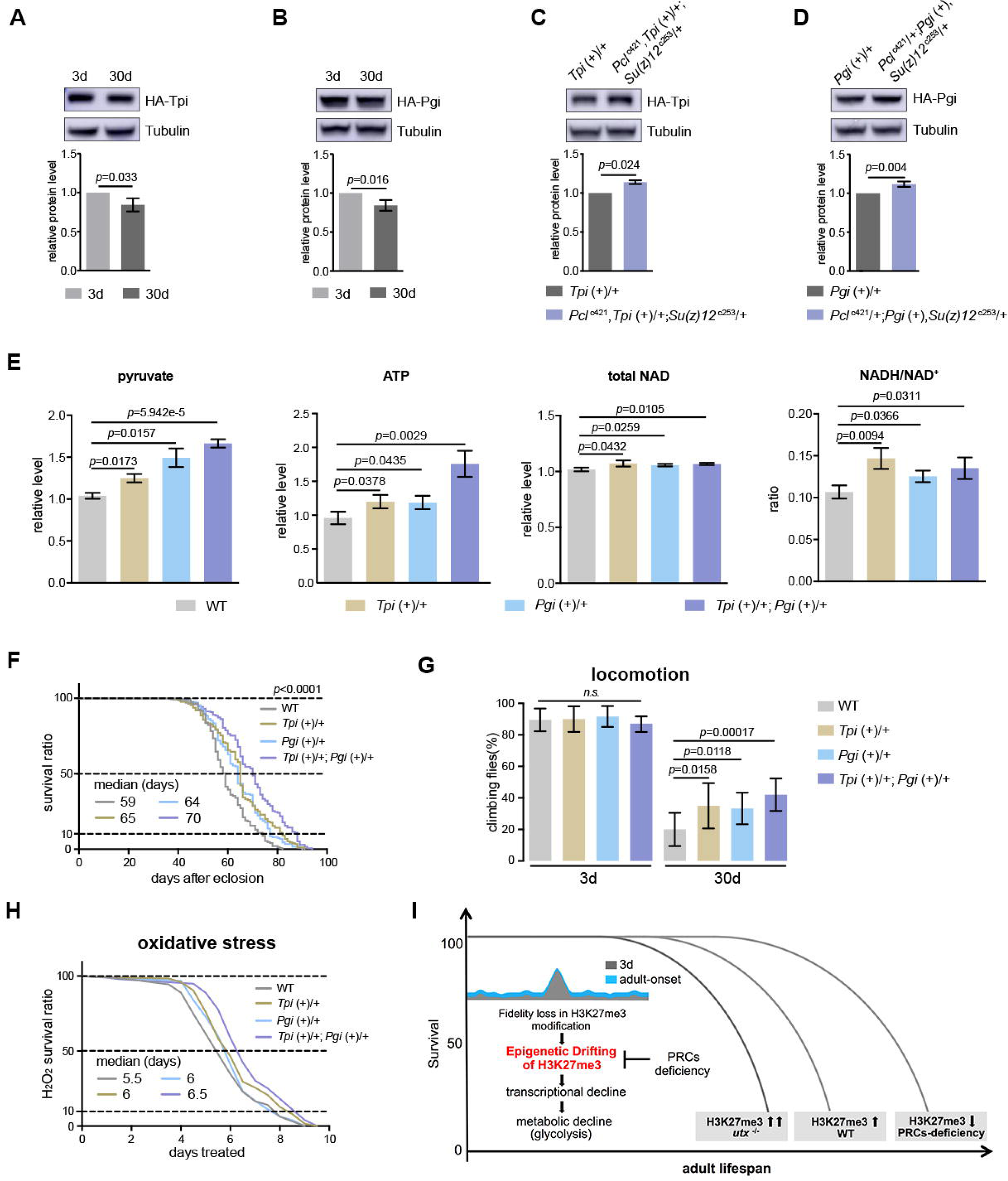
Transgenic increase of glycolytic genes suffices to elevate glycolysis and healthy lifespan. (A) Tpi protein western blot (top) and quantification (bottom) shows a decrease with age. (mean±SD of 3 biological repeats; student t-test). Western blot was from 3d and 30d old male flies. Genotype: *Tpi* (+)/+. (B) Pgi protein western blot (top) and quantification (bottom) shows a decrease with age. Statistics and Western blot as in (A). Genotype: *Pgi* (+)/+. (C) Tpi protein western blot (top) and quantification (bottom) shows an increase in PRC2 mutants. Statistics and Western blot as in (A). Genotype: *Tpi* (+)/+. *Pcl*^c421^, *Tpi* (+)/+; *Su(z)12*^c253^/+. (D) Pgi protein western blot (top) and quantification (bottom) shows an increase in PRC2 mutants. Statistics and Western blot as in (A). Genotype: *Pgi* (+)/+. *Pcl*^c421^/+; *Pgi* (+), *Su(z)12*^c253^/+. (E) Transgenic increase of glycolytic genes stimulates glycolysis, as shown by elevated pyruvate, ATP, and NADH/NAD^+^ ratio compared to age-matched WT (mean±SD of 3 biological repeats; student *t*-test). Metabolite analysis was from muscle tissues of 30d old male flies. Genotypes: WT: 5905. *Tpi* (+)/+. *Pgi* (+)/+. *Tpi* (+)/+; *Pgi* (+)/+ (F-H) Transgenic increase of glycolytic genes promotes adult fitness, including lifespan (F), locomotion (G), and resistance to oxidative stress (H). (for lifespan assay: 25°C; n≥200 per genotype for curve; log-rank test; for climbing assay: mean±SD of 10 biological repeats with 10 flies for each repeat; student *t*-test; for oxidation assay: 25°C; n=100 per genotype for curve; log-rank test). Genotypes as in (E). (I) Model. Adult-onset fidelity loss results in epigenetic drift of H3K27me3. Over a chronic timescale, the drifting of H3K27me3 induces transcriptional and metabolic decline including a reduction of glycolysis. Effects of PRC2-deficiency in life-extension can be at least in part attributed to the effect in stimulation of glycolysis, thereby maintaining metabolic health and longevity. Adult lifespan is inherently modulated by the alterations in the levels of H3K27me3, as shown by the corresponding change during natural aging, in *utx* mutants as well as in PRCs-deficiency.

Taken together, these data suggest that upregulation of glycolytic genes alone recapitulates anti-aging features of PRC2 mutants. This finding, combined with data obtained from H3K27me3 dynamics and consequently its effects on gene expression and metabolism, underscores the mechanistic link between epigenetic, transcriptional, and metabolic processes in aging, further heightening the role of glycolysis in promotion of metabolic health and longevity.

## DISCUSSION

Whether epigenetic changes during adult lifespan is merely associated with or directly contribute to it is a long-standing question in the biology of aging. Our study provides a new paradigm by which epigenetic drift of H3K27me3, a highly conserved histone mark, links glycolysis to healthy longevity (Figure 7I). Though H3K27me3 levels have been shown to increase with age in mammals (Liu et al., 2013; Sun et al., 2014) and flies (current study), the biological significance of such changes on aging is unknown. Our quantitative assessment shows that aging leads to a decline of fidelity in H3K27me3 modification, as demonstrated by a genome-wide drift of this repressive mark. As a consequence, glycolytic genes are downregulated, which in turn causes a reduction of glycolysis and glucose metabolism. Importantly, the age-delaying effect of PRCs-mutation likely arises from its ability to slow down or reverse this trend. Therefore we suggest that epigenetic drift of H3K27me3, enhanced by its age-modulated increase, is a new mechanism that drives the progression of aging. Expression of *Tpi* and *Pgi* of glycolytic genes is inherently regulated by the alterations of H3K27me3 levels, as shown by their corresponding changes during natural aging, in *utx* mutants as well as in PRCs-deficiency. We propose that H3K27me3 dynamics may elicit changes in local chromatin environment—especially inter-peak regions that are potentially more sensitive to the gain or loss of the epigenetic mark—thus enabling fine-tuning of target gene expression.

It is unclear how aging might promote the drift in H3K27me3 modification. One possible reason is age-associated DNA damage that reduces the fidelity of epigenetic markings. It has been well-established that aging is associated with the accumulation of DNA damage (Lu et al., 2004). In *Bombyx mori,* UV-C irradiation can induce the increase of H3K27me3 mediated by PRC2 (Li et al., 2014). In *Neurospora crassa,* on the other hand, the increase and redistribution of H3K27me3 can be induced by the loss of H3K9 methyltransferase complex (Basenko et al., 2015). Thus, dysregulation of H3K27me3 may be due to feed-forward interactions between DNA damage and changes in the levels of complexes that control epigenetic markings during aging.

Metabolic control has emerged as a critical player to determine the aging outcome. We propose that stimulation of glycolysis promotes healthy longevity. Previous data implicates that impairing glycolysis can extend lifespan perhaps by triggering compensatory processes (Schulz et al., 2007). However, rat models supplemented with 2-deoxyglucose, a known anti-glycolytic compound, are instead short-lived (Minor et al., 2010). Moreover, glucose-enriched diet can prevent pathophysiological decline and early death of telomere dysfunctional mice by stimulating glucose metabolism including glycolysis (Missios et al., 2014). Importantly, recent evidence from studies on a wide range of species suggests that a balanced diet from protein to carbohydrate and their interactive effects are key elements for healthy longevity. Mouse dietary studies have demonstrated that animals fed on diets that are low in protein and high in carbohydrate enjoy longer life with lower blood pressure, improved glucose tolerance and lipid profiles (Solon-Biet et al., 2014). This is also supported by human data on the deleterious effects of diets with high protein and low-carbohydrate (Floegel and Pischon, 2012; Lagiou et al., 2012). Hence the impact of glucose metabolism including glycolysis on aging might be poised to integrate age-associated physiology; the precise outcome must be rather complicated, which could be influenced not only by diverse spatiotemporal need in normal or pathological conditions, but also upon newly evolved biological activities correlated with the increased complexity from nematode to *Drosophila* and mammals. Our findings propose that the ability to stimulate glycolysis is critical to ensure a healthy level of physical capability by delaying the bioenergetic decline during aging as which otherwise might form a vicious cycle from decrease in physical activity to hypometabolism, and perhaps disease susceptibility.

Increased ratios of GSH/(GSH+GSSG) and NADH/NAD^+^ produced from enhanced glycolytic activities may provide a simple but effective way to retard aging. The oxidative phosphorylation, though produces more ATP than glycolysis, can yield intracellular ROS. The accumulation of ROS is the leading proposed cause of decline in cellular function and integrity in aging (Balaban et al., 2005). Thus, modulating H3K27me3 may reprogram bioenergetic decline during aging, which in effect reduces cellular damage and deterioration. Importantly, mammalian glycolytic genes have also been shown as PRCs targets (Brookes et al., 2012). Future investigations, including in-depth comparative analysis of PRCs and glycolytic pathway in the aging process in both flies and humans, may harness common operative mechanisms that modulate metabolic homeostasis and healthy longevity. Given the reversible nature of epigenetic pathways, this study proffers a tempting strategy against age-associated physiological decline and disease.

## AUTHOR CONTRIBUTIONS

Conceptualization, N.L.; Methodology, Hui.W., Y.C., and Z.Z.; Investigation, Z.M., Hui.W., Y.C., Han.W., K.N., X.W., H.M., Y.Y., and W.T.; Data Curation, Hui.W., Y.C., Z.L., F.L., Y.Z., R.L., Z.Z., and N.L.; Wring – Original Draft, Hui.W. and N.L.; Writing – Review & Editing, Z.M., Hui.W., and N.L.; Funding Acquisition, Z.L., Y.Z., Z.Z., and N.L.; Supervision, N.L.

## ACKNOWLEDGEMENTS

We thank Drs. Jilong Liu, Yun Zhao, Meng-Qiu Dong, Cong Liu, Chao Tong, Xing Guo, Lanfeng Wang, and Ye Tian for advice on the manuscript; Wei Wu and the Core Facility of *Drosophila* Resource and Technology, Shanghai Institute of Biological Sciences, Chinese Academy of Sciences, Shanghai, China for fly microinjections. Prof. Junying Yuan provided considerable support, advice on the experiments, and critical suggestions on the manuscript. N.L. and Z.Z. are Junior Scholar of the 1000 Plan of China. Y.Z. is supported by 100 Talents Program of the Chinese Academy of Sciences. This work was supported by grants from the National Program on Key Basic Research Project of China to N.L. and Y.Z. (2016YFA0501900); the National Natural Science Foundation of China to N.L. (31371326), Y.Z. (31671428, 31500665, 31530041), Z.Z. (21575151), and F.L. (81770143); a National Institute of Health grant (GM120033) to Z.L.; a National Science Foundation grant (DMS-1263932) to Z.L.; a Cancer Prevention Research Institute of Texas grant (RP170387) to Z.L.; the Robert A. and Renée E. Belfer Family Foundation.to Z.L., and the Chao Family Foundation to Z.L.

---

**Figure S1. Fidelity loss in H3K27me3 modification results in its genome-wide drifting.**

(A) Western blot (top) and quantification (bottom) show an increase of H3K27me3 with age in head (left) and muscle (right) tissues. (mean±SD of 3 biological repeats; student *t*-test). Genotype: 5905.

(B) Quantitative ChIP-Rx protocol. A fixed amount of the mouse epigenome was introduced as the spike-in reference, thereby allowing quantitative and comparative assessment of the fly epigenomes.

(C) Venn diagram shows common peak regions from four replicates. Number of peaks and genomic coverage were indicated in bold black. ChIP-seq was from muscle tissues of 3d old male flies. Genotype as in (A).

(D) Pairwise Pearson correlation analysis (left) and venn diagram (right) between 3d and 30d ChIP-seq datasets of WT. Four replicates for each age group (rep1-4). Peaks reproducibly identified by replicates were used to make the venn diagram. ChIP-seq was from muscle tissues of 3d and 30d old male flies. Genotype as in (A).

**Figure S2. Long-lived PRC2 mutants diminish the epigenetic drift of H3K27me3 during aging.**

(A) H3K27me3 western blot (left), H3K27me3 quantification (middle), and lifespan curve (right) for pair-wise combinations of PRC2 trans-heterozygous double mutants. (for H3K27me3 quantification: mean±SD of 3 biological repeats; student t-test; for lifespan assay:
25°C; n≥200 per genotype for curve; log-rank test). Western blot was from head tissues of 3d old male flies.

(B) Relative quantification of different histone marks between WT and PRC2 mutants. (for quantification: mean±SD of 3 biological repeats; student *t*-test; ***p<0.001). Western blot as in (A). Genotypes: WT: 5905. Mut: *Pcl*^c421^/+; *Su(z)12*^c253^/+.

(C) and (D) Pairwise Pearson correlation analysis and venn diagram between WT and PRC2 mutants. ChIP-seq was from muscle tissues of 3d old male flies. Two replicates for each genotype (rep1-2). Peaks reproducibly identified by replicates were used to make the venn diagram. Genotypes: WT: 5905. *Pcl*^c421^/+; *Su(z)12*^c253^/+. *esc*^c289^/+; *E(z)*^c239^/+.

(E) H3K27me3 modification decreases in PRC2 mutants. Scatter plot showed H3K27me3 signals of 17511 genes in *esc:*^289^; *E(z)*^239^ as compared to WT. (wilcoxon signed-rank test, *p*<2.2e-16). X- and Y-axis represented signal intensity transformed by Log2. Contour lines indicated that H3K27me3 signals were higher in WT compared to mutants. ChIP-seq was from muscle tissues of 3d old male flies. Genotypes: WT: 5905. *escc^c289^/+; *E(z)*^c239^/+.*

(F) Inter-peak genes receive relatively less H3K27me3 modification in PRC2 mutants. Violin plots for inter-peak genes (left) and peak genes (right) between WT and mutants. We computed bootstrapped confidence intervals for the mean ChIP intensity (10,000 draws with replacement of n=500). Net loss of H3K27me3 signals was calculated by a subtraction of the mean signal intensity between WT and mutants. Bootstrapped 95% confidence intervals: WT inter-peak genes: [2.626, 3.601]; *esc^289^; E(z)^239^* inter-peak genes: [2.029, 2.627]; WT peak genes: [9.161, 10.367]; *esc^289^; E(z)^239^*inter-peak genes: [7.363, 8.355]. ChIP-seq was from muscle tissues of 3d old male flies. Genotypes: WT: 5905. *escc*^c289^/+; *E(z)*^c239^/+.

(G) Modification of H3K27me3 during aging has reduced selectivity in PRC2 mutants. Bootstrapped 95% confidence intervals: *esc^289^; E(z)*^239^ late-onset inter-peak genes: [0.551, 0.804]; *esc^289^; E(z)^239^* late-onset peak genes: [0.532, 0.845]. ChIP-seq was from muscle tissues of 3d and 30d old male flies. Genotypes: *escc*^c289^/+; *E(z)*^c239^/+.

(H) Age-associated drifting of H3K27me3 is dampened in PRC2 mutants. Scatter plot for 17511 genes between WT and mutants. X- and Y-axis represented signal intensity transformed by Log2. Contour lines indicated that the majority of the gene signals displayed higher levels—thus more rapid changes—in aged WT compared to mutants. ChIP-seq and genotypes as in (F).

**Figure S3. Transcriptomics links H3K27me3 dynamics to the regulation of glycolytic genes.**

(A) Number of genes upregulated in PRC2 mutants with indicated genotype (cutoff: *p*<0.05). Note that transcriptional changes were generally mild, and the majority of changes were genes in the inter-peak regions.

(B) Diagram illustrates that the transcriptional changes of genes mediating glucose and the citric acid cycle. Genes and their relative transcriptional changes in 3d and 30d old WT and PRC2 long-lived mutants with indicated genotype (cutoff *p*<0.05; *n.s.:* not significant). Commonly changed genes, *Tpi* and *Pgi*, were highlighted in red. While anaerobic glycolysis yields ATP only, glycolysis at the step of pyruvate formation has a net outcome of both ATP and NADH.

(C) ChIP-qPCR of glycolytic genes *Tpi* (top panel) and *Pgi* (bottom panel). Genomic structure and gene isoforms were shown according to the Flybase annotation. Along the gene of indicated genotype, a, b, c, and d were underlined, denoting the sites for PCR amplification. Color codes represented ATG (the translation start site), CDS (coding sequence), 5’UTR, and 3’UTR. (mean±SD of 3 biological repeats; student *t*-test). ChIP-qPCR was from muscle tissues of 3d and 30d old male flies. Genotypes: WT: 5905. *Pcl*^c421^/+; *Su(z)12*^c253^/+. *utx*^c499/c499^.

(D) and (E) Short-lived *miR-34* mutants (D) and long-lived *piwi* mutants (E) demonstrated that changes of the glycolytic genes had no simple correlation with lifespan alterations. (for lifespan assay: 25°C; n≥200 per genotype for curve; log-rank test). Genotypes: WT: 5905. *miR-34*^c120/c120^. *piw*^c362^/+.

**Figure S4. Untargeted metabolomics.**

(A) Heatmap of metabolites exhibiting differential levels of 3d, 30d WT and 30d PRC2 mutants. Metabolomics was from head tissues of 3d and 30d old male flies. Genotypes: WT: 5905. *Pcl*^c421^/+; *Su(z)*^c253^/+.

(B) Metabolomic analysis reveals that metabolites of the citric acid cycle show comparable levels between young and aged WT, and between WT and PRC2 mutants. *(n.s.:* not significant). Metabolomics and genotypes as in (A).

**Figure S5. Metabolic glucose flux experiment.**

(A) Schematic representation of the glucose flux experiment in adult aging flies. Briefly, flies at 3d (young) and 25d (aged) were switched from ^12^C-glucose to ^13^C-glucose food for a fixed five days, such that the glucose metabolism can be traced by measuring the ^13^C-labeled metabolites.

(B) Relative to the total glucose pool, ^13^C-glucose accounts for more than 93% and 95% in 8d and 30d old animals, respectively, suggesting a near-complete replacement. (*n.s.:* not significant). Metabolomics was from head tissues of 8d and 30d old male flies. Genotypes: WT: 5905. *Pcl*^c421^/+; *Su(z)12*^c253^/+.

(C) Metabolic flux analysis reveals that the levels of ^13^C_6_-labeled glucose are comparable between WT and mutants. (mean±SD of 8 biological repeats; wilcoxon rank-sum test). Metabolomics and genotypes as in (B).

(D) and (E) Quantification of isotopologues pertaining to each metabolites of citric acid cycle (D) and their summed intensity (E) indicate no consistent change of the pathway with age and in PRC2-deficiecny. Metabolomics and genotypes as in (B).

**Figure S6. *Tpi* and *Pgi* deficient flies have normal adult phenotypes.**

Assessments of adult lifespan (A) and (E), oxidation test (B) and (F), climbing ability (C) and (G), and the levels of ATP and the ratio of NADH/NAD^+^ (D) and (H), show no difference between WT and flies with either *Tpi* or *Pgi* heterozygous mutation. (for lifespan assay: 25°C; n>190 per genotype for curve; log-rank test; for oxidation tests: 25°C; n=100 per genotype for curve; log-rank test; for climbing assay: mean±SD of 10 biological for each repeats; student *t*-test; for analysis of metabolites: mean±SD of 3 biological for each repeats; student *t*-test; *n.s.:*not significant). Genotypes: WT: 5905. *Tpf*^c511^/+. *Pgi*^c392^/+.

**Figure S7. Genomic transgenes for *Tpi* and *Pgi*.**

(A) and (B) Genomic structure and gene isoforms were shown according to the Flybase annotation. Color codes represented ATG (the translation start site), HA tag, CDS (coding sequence), 5’UTR, and 3’UTR. Positive control was based on the UAS/Gal4 system and overexpression of cDNAs for *Tpi* and *Pgi,* respectively. Transient transfection of *Tpi* and *Pgi*genomic constructs yielded protein of appropriate size. Western blot was using *Drosophila*S2R^+^ cells.

**Table S1. Summary of H3K27me3 peak regions.**

Summary of ChIP-seq experiments. Two criteria for peak calling: IP/input≥2, signals spanning ≥3kb. In total, 222 peak regions of H3K27me3 were reproducibly identified by four biological replicates at 3d. For each peak regions, information pertaining to chromosomal location and genes therein contained were given. ChIP-seq was using muscle tissues of 3d old animals. Genotype: 5905.

**Table S2. Summary of CRISPR/Cas9-led gene mutagenesis and lifespan.**

For each gene, two sgRNAs with indicated sequences, detailed deletion sites confirmed by Sanger sequencing, and PCR validation were shown. To name new CRISPR mutant, a superscript amended to the gene contained a letter c denoting CRISPR/Cas9-directed mutagenesis followed by the size of genomic deletion. Mutants were backcrossed (at least 5 generations) into 5905 (Flybase ID FBst0005905, w^1118^) to assure that phenotypes were not associated with any variation in background. Lifespan curves, together with 50% (median lifespan) and 10% survival data were listed. (for lifespan assay: 25°C; n≥140 per genotype for curve; log-rank test).

**Table S3. Summary of transcriptomic analysis of PRC2 longevity mutants.**

RNA-seq summary. This table summarizes 5 categories of genes and their transcriptional changes in long-lived PRC2 mutants of indicated genotype: (1) commonly upregulated and (2) downregulated genes in all PRC2 long-lived mutants; genes known to promote longevity upon upregulation, including (3) insulin pathway genes, (4) TOR signaling pathway genes, and (5) genes of miscellaneous pathways. RNA-seq was from muscle tissues of 30d old male flies.

**Table S4. Summary of transcriptomic analysis of genes in glycolysis, citric acid cycle, pentose phosphorylation pathway, and oxidative phosphorylation.**

This table summarizes metabolic genes and their transcriptional changes in long-lived PRC2 mutants of indicated genotype. RNA-seq was from head and muscle tissues of 3d and 30d old male flies.

**Table S5. Primer sequences used for the study.**

## EXPERIMENTAL MODEL AND SUBJECT DETAILS

### Fly culture

Flies were cultured in standard media at 25°C with 60% humidity in a 12h light and 12 h dark cycle unless otherwise specified. The WT line used was 5905 (FlyBase ID FBst0005905, *w^1118^*). All fly lines used in this study have been backcrossed with 5905 for five consecutive generations for a uniform genetic background, to assure that phenotypes were not associated with any variation in background.

## METHOD DETAILS

### Western blot assays for different histone modifications

50 heads or 30 muscles were harvested and immediately frozen in liquid nitrogen. Tissues were homogenized by Dounce Homogenizer. Two different methods were used for histone extraction: nuclei isolation and total protein extract treated by sonication. For nuclei isolation, Aogma nucleus-cytoplasm extraction kit was used (Aogma). For total protein extract, tissues were lysed by RIPA buffer and then treated with 5 cycles of sonication with 30 seconds on and 30 seconds off (Bioruptor Pico). Resulted histone extractions were loaded onto a NuPAGE 12% Bis-Tris Gel (Thermo Fisher Scientific), and then transferred by electrophoresis to a polyvinylidene fluoride membrane (Millipore). The corresponding primary antibodies were diluted 1:1000 and incubated with the membranes at 4°C overnight, the HRP-labeled secondary antibodies were diluted 1:40000 and incubated for 1 hour at room temperature. The signals were detected using ECL (Thermo Fisher Scientific) by Amersham Imager 600 (GE Healthcare). The signal intensity was quantified by ImageQuant TL (GE Healthcare). The relative normalization of histone modifications was normalized to the histone H3 signal. Three biological replicates were done for each experiment.

### CRIPSR/Cas9 mutagenesis

CRISPR/Cas9 mutagenesis was performed as previously described (Ren et al., 2013a). Two sgRNA plasmids for target gene were injected into fly embryo. Single-fly-PCR assays were used to screen mutants. To do this, single fly was homogenized in 50μl squashing buffer (10 mM Tris buffer (pH 8.5), 25 mM NaCl, 1 mM EDTA, 200 μg/ml Proteinase K), then incubated at 37°C for 30 minutes, followed by 95°C, 10 minutes for inactivation. Screen primers for each target genes were listed in Table S2. The virgin females carrying the deletion were backcrossed into WT (5905) male flies for five consecutive generations for a uniform genetic background, to mitigate background effects.

### Genomic DNA transgene

Genomic DNA covering the Tpi and Pgi genes was prepared by PCR-amplification and fusion PCR. DNA was clone into the pBID vector (Wang et al., 2012). Fly microinjection and transgenesis followed standard procedures.

### Metabolic flux analysis using U-[^13^C]-glucose

160 male flies for each genotype at 3d or 25d of age were used, with 20 flies per vial. Prior to the test, flies were pre-treated with 1% Agar media for 6 hours before transferred to the vials containing a small piece of Kimwipe filter paper pre-soaked in 1 ml of 10% U-^13^C_6_-glucose. Flies were treated for 3 days, and then transferred to new vials with fresh U-^13^C_6_-glucose for additional 2 days. Fly heads were dissected for subsequent metabolic analysis. For each genotype, 8 biological repeats were conducted, with 20 heads for each repeats.

### Assays for adult phenotypes

#### Lifespan assays

Adult male flies were collected at the day of eclosion and maintained at 20 flies per vial at 25°C with 60% humidity and a 12h light/12h dark cycle. Flies were transferred to new vials every other day and scored for survival. The Prism software (Graphpad) was used to draw the survival curve and statistical analysis.

#### Climbing ability assay

Ten male flies were transferred to an empty vial for 30 minutes adaption in dark. Flies were tapped to the bottom of the vial; the percentage of flies failed to climb up to a mark at 2 cm from the bottom within 5 seconds was scored. Ten biological replicates were done for each genotype at given age.

#### Oxidative stress

100 male flies for each genotype at given age were used, with 20 flies per vial. Prior to the test, flies were pre-treated with 1% Agar media for 6 hours before transferred to the media containing oxidative radical species. For H_2_O_2_ treatment, flies were transferred to vials containing a small piece of Kimwipe filter paper pre-soaked in 1.5 ml of 10% glucose and 2% H_2_O_2_ (Sinopharm Chemical Reagent). Dead flies were scored every 12 hours, and the Prism software (Graphpad) was used to generate the survival curve and statistical analysis.

### Sequencing experiments

#### ChIP-seq

For developmental stages, embryo (30min-1 hour), larva (96 hour), and pupae (7d) were analyzed. For adult stage, 100 fly muscles at each time point were used. Samples were collected into a 1.5 ml microcentrifuge tube and immediately frozen in liquid nitrogen. Tissues were pulverized to fine powder using a liquid nitrogen cooled mortar and pestle. Resulted fine power was resuspended in 1.2 ml PBS; 32.4 μl of 37% formaldehyde was added for cross-linking at room temperature for 10 minutes. To quench formaldehyde, 60 μl of 2.5 M glycine was added. Centrifugation was using 5,000 g for 5 minutes at 4°C. After removing the supernatant, pellet was washed twice with 1 ml PBS containing proteinase inhibitor (Roche). This lysis step was performed at 4°C for more than 1 hour in RIPA buffer (Sigma) supplemented with proteinase inhibitor. The lysate was sonicated using Bioruptor Pico (Diagenode), for 15 cycles with 30 seconds on and 30 seconds off. Centrifugation was using 10,000g for 10 minutes at 4°C. Supernatant was carefully transferred to a new tube without disturbing the debris. Pre-clear was performed using ChIP-Grade Protein A/G Plus Agarose (Thermo Fisher Scientific) at 4°C for 1 hour. For spike-in epigenome, mouse neuro-2a cells were prepared with the same procedures. Prior to mixing mouse epigenome with that of fly’s, 5% from each preparations was transferred for quantification of DNA mass. To do this, RIPA buffer was added to a final volume of 120 μl, followed by 1 μl RNase A to remove the RNA at 37°C for 30 minutes. Samples were treated with 0.2 M NaCl (final concentration) at 65°C for 4 hours and 200 ng/μl proteinase K (final concentration) at 55°C for 2 hours. DNA was purified by PCR purification kit (Qiagen), and DNA mass was quantified by Qubit dsDNA HS assay kit (Thermo Fisher Scientific). According to the DNA mass, 5% of the mouse epigenome was added to the fly sample. 10% of sample was transferred to a new tube as ChIP input control. Immunoprecipitation was performed by incubating 3 μg antibody at 4°C for 5 hours with rotation. ChIP-Grade Protein A/G Plus Agarose was added. The sample was incubated overnight at 4°C with rotation. Beads were washed twice with ChIP Wash Buffer (150mM NaCl, 2mM EDTA (pH8.0), 20mM Tris- HCl (pH8.0), 1% Triton X-100, and 0.1% SDS), and another two washes using ChIP Final Wash Buffer (500mM NaCl, 2mM EDTA (pH8.0), 20mM Tris-HCl (pH8.0), 1% Triton X-100, and 0.1% SDS). The elution step was using 120 μl of ChIP Elution Buffer (100mM NaHCO_3_ and 1% SDS) by incubating at 65°C for 30 minutes. Resulted DNA was purified as above described, followed by RNase A treatment. 5-10 ng of DNA was used to generate sequencing library using NEB DNA library prep kit. The quality of libraries was checked by Bioanalyzer 2100 (Agilent). The quantification was performed by qRT-PCR with a reference to a standard library. The libraries were pooled together in equimolar amounts to a final 2 nM concentration. The normalized libraries were denatured with 0.1 M NaOH (Sigma). Pooled libraries were sequenced on the illumina Miseq/Next-seq platform with single end 100 bps.

#### polyA-selected RNA-seq

Thirty muscles were dissected and homogenized in a 1.5 ml tube containing 1 ml of Trizol Reagent (Thermo Fisher Scientific). RNA isolation was followed in accordance with manufacturers instruction. RNA was resuspended in DEPC-treated RNase free water (Thermo Fisher Scientific). TURBO DNA free kit was used to remove residual DNA contamination according to manufacturers instruction (Thermo Fisher Scientific). 1 μg of total RNA was used for sequencing library preparation. PolyA-tailed RNAs were selected by NEBNext Poly(A) mRNA Magnetic Isolation Module (NEB), followed by the library prep using NEBNext Ultra RNA library Prep Kit for Illumina according to manufacturers instruction (NEB). The libraries were checked and pooled as described above. Pooled denature libraries were sequenced on the illumina NextSeq 550 or Hiseq 2500 platforms with single end 100 bps.

### Quantitative PCR

#### ChIP-qPCR

ChIP experiments were performed as described above. The same volume of input and IP DNA were used as template for quantitative PCR experiment. Analysis was performed using the QuantStudio 6 Flex real-time PCR system with SYBR selected master mix (Thermo Fisher Scientific). The percentage of input method was used for data analysis. Primers for target genes were listed in Table S5.

#### RT-qPCR

RNA extractions were done as described above. 1 μg of total RNA was used for reverse transcription by random primers using SuperScript III First-strand synthesis system for RT-PCR (Thermo Fisher Scientific). Analysis was performed using the QuantStudio 6 Flex real-time PCR system with SYBR selected master mix (Thermo Fisher Scientific). The 2^-ΔΔCT^ method was used for quantification upon normalization to the RP49 gene as internal control. Primers for target genes were listed in Table S5.

### Metabolome analysis

#### Metabolites extraction

20 heads for each replicates were collected into a microcentrifuge tube and immediately frozen into liquid nitrogen. For each experiment, 10 biological replicates were prepared. Fly heads were homogenized with 200 μl of H_2_O and 20 ceramic beads (0.1 mm) using the homogenizer (Precellys 24, Bertin Technologies). 800 μl of ACN:MeOH (1:1, v/v) was added for metabolite extraction. Vortex was for 30s, followed by incubation in liquid nitrogen for 1 min. This freeze-thaw cycle was repeated three times. To precipitate protein, samples were incubated for 1 h at −20 °C, followed by 15 min centrifugation using 15,000g at 4 °C. The supernatant was removed and evaporated to dryness in a vacuum concentrator (Labconco, German). Dry extracts were then reconstituted in 100 μl of ACN:H_2_O (1:1, v/v), followed by 10 min sonication (50Hz, 4°C) and 15 min centrifugation using 13,000 rpm at 4 °C to remove insoluble debris. Supernatants were transferred to HPLC glass vials and stored at −80 °C prior to LC/MS analysis.

#### LC-MS/MS

LC-MS was performed using a UHPLC system (1290 series, Agilent Technologies) coupled to a quadruple time-of-flight mass spectrometer (AB Sciex TripleTOF 6600). Waters ACQUITY UPLC BEH Amide columns (particle size, 1.7μm; 100 mm (length) × 2.1 mm (i.d.)) were used. Mobile phases A=25 mM ammonium acetate and 25 mM ammonium hydroxide in 100% water, and B=100% acetonitrile, were used for both ESI positive and negative modes. And the linear gradient eluted from 95 % B (0.0-1.0 min), 95% B to 65% B (1.0-14.0 min), 65% B to 40% B (14.0-16.0 min), 40% B (16.0-18.0 min), 40% B to 95 % B (18.0-18.1 min), then stayed at 95% B for 4.9 minutes. The flow rate was 0.3 mL/min and the sample injection volume was 2 μl.

ESI source parameters on TripleTOF 6600 were set as followings: ion source gas 1 (GS1), 60 psi; ion source Gas 2 (GS2), 60 psi; curtain gas (CUR), 30 psi; temperature (TEM), 600 °C; ion spray Voltage floating (ISVF), 5000 V or −4000 V, in positive or negative modes, respectively; declustering potential (DP), 60 V or −60 V in positive or negative modes, and collision energy for Product Ion, 30 eV or −30 eV, in positive or negative modes, respectively; collision energy spread (CES), 0 eV. LC-MS/MS data acquisition was operated under information-dependent acquisition (IDA) mode. The instrument was set to acquire over the m/z range 60-1200 Da for TOF MS scan and the m/z range 25-1200 for product ion scan. The accumulation time for TOF MS scan was set at 0.20 s/spectra and product ion scan at 0.05 s/spectra. The unit resolution was selected for precursor ion selection, and the collision energy (CE) was fixed at 30 V and −30 V for positive or negative modes, respectively. Collision energy spread (CES) was set as 0. IDA settings which were set as followings: charge state 1 to 1, intensity 100 cps, exclude isotopes within 4 Da, mass tolerance 10 ppm and maximum number of candidate ions 6. The “exclude former target ions” was set as 4 seconds after 2 occurrences. In IDA Advanced tab, “dynamic background subtract” was also chosen.

### Qualitative metabolic flux analysis using U-[^13^C]-glucose as a tracer

#### Metabolites extraction

Metabolites were extracted as described above for untargeted metabolomics.

#### LC-MS

LC-MS analysis was performed using a UHPLC system (1290 series, Agilent Technologies) coupled to a quadruple time-of-flight mass spectrometer (6550 series, Agilent Technologies). Merck SeQuant ZIC-pHILIC column (particle size, 5 μm; 100 mm (length) × 2.1 mm (i.d.)) was used. Mobile phases A=25 mM ammonium acetate and 25 mM ammonium hydroxide in 100% water, and B=100% acetonitrile, were used for both ESI positive and negative modes. And the linear gradient eluted from 80 % B (0.0-2.0 min, 0.2 mL/min), 80% B to 20% B (2.0-17.0 min, 0.2 mL/min), 20% B to 80% B (17.0-17.1 min, 0.2 mL/min), 80% B (17.1-22.1 min, 0.4 mL/min), 80% B to 80 % B (22.1-22.2 min, 0.2 mL/min). The sample injection volume was 2 μL.

ESI source parameters on QTOF 6550 were set as followings: sheath gas temperature, 300 °C; dry gas temperature, 250 °C; sheath gas flow, 12 L/min; dry gas flow, 16 L/min; capillary voltage, 2500 V or −2500 V in positive or negative modes, respectively; nozzle voltage, 0 V; and nebulizer pressure, 20 psi. The MS1 data acquisition frequency was set as 4 Hz, and the TOF scan range was set as m/z 60 - 1200 Da.

#### Data analysis

Data processing for qualitative metabolic flux analysis was performed using VistaFlux software suite from Agilent Technologies. First, Pathways to PCDL software (version B.07.00, Agilent Technologies) and PCDL Manager software (version B.07.00, Agilent Technologies) was used to build a metabolite library for metabolites in both glycolysis and citric acid cycle. Then, acquired LC-MS raw data files (.d) were loaded into Profinder (version B.08.00, Agilent Technologies) for the extraction of metabolite isotopologues using the constructed metabolite library. The ion abundance criterion for data extraction was set as “use peak core area 20% of peak height”. The qualification parameters were set as: mass tolerance: ± 15 ppm + 2.00 mDa; retention time tolerance: ± 0.20 minutes; anchor ion height threshold: 250 counts; sum of ion heights threshold: 1000 counts; correlation coefficient threshold: 0.5. The final results were exported into a csv file for the subsequent comparative and statistical analyses.

#### Specific metabolites tests

Ten fly muscles were harvested and immediately frozen in liquid nitrogen. Tissues were homogenized using the homogenizer (Precellys 24, Bertin Technologies) with 300 μl ddH_2_O, or extraction buffer provided by the kit. For glucose, ATP, GSH and pyruvate assays, tissues were homogenized with ddH_2_O, then frozen at −80°C for 30 minutes and thawed at room temperature, followed by 5 cycles of 30 seconds on and 30 seconds off sonication. For NADH and NADPH assays, tissues were homogenized with extraction buffer provided by the kit. Lysates were centrifuged using 10,000 g at 4°C for 5 minutes and the supernatant was transferred to 10 kDa molecular weight cut-off spin filter (Sigma), followed by centrifugation using 10,000 g at 4°C for 30 minutes for deproteinization. Assays were assembled according to the instruction of manufacturers (Glucose [Sigma, MAHK20], ATP [Promega, FF2000], GSH [Abcam, ab13881], pyruvate [Sigma, MAK071], NADH [Sigma, MAK037], NADPH [Sigma, MAK038]). Signals were detected by the EnSpire Multimode Plate Reader (PerKin Elmer).

### Bioinformatic analyses

#### ChIP-seq

Sequencing reads were mapped to the reference genome dm6 or mm10 respectively with Bowtie2-2.2.9 (Langmead and Salzberg, 2012) by default parameters. Samtools-1.3.1 (Li et al., 2009) was used for sam to bam format conversion. Peak regions were identified by homer-v4.8.3 (Heinz et al., 2010) function findPeaks with parameter “-style histone -F 2 -size 3000 -minDist 5000”. The overlapped peaks of replicate datasets were found using homer function mergePeaks by default parameter. The intersecting peak regions between different genotypes and aging stages were identified by BEDTools-2.26.0 (Quinlan and Hall, 2010) by default parameter. The Pearson correlation numbers were obtained by Galaxy/Cistrome-1.0.0 (Liu et al., 2011) and compared at a step size of 1000 bps. For quantitative comparison, the dm6 mapped reads were normalized to a scale factor using deeptools-2.2.4 (Ramirez et al., 2014) function bamCoverage with 10 bp bin size. The scale factor for each sample was calculated as reported (Orlando et al., 2014) with a small modification. The percentage of mouse genome in input was set to the mm10 mapped reads in total mapping reads, rather than a constant. The ChIP intensity of each gene or region was calculated using the Bwtool (Pohl and Beato, 2014) function bwtool summary with default parameters. The Circos plot was generated by J-circos-V1 (An et al., 2015). The selected genome regions were displayed by IGV-2.3.31 (Robinson et al., 2011) with the bigwig files generated by bamCoverage.

#### PolyA-selected RNA-seq

Sequencing reads were mapped to the reference genome dm6 with STAR2.3.0e (Dobin et al., 2013) by default parameter. The read counts for each gene were calculated by HTSeq-0.5.4e (Anders et al., 2015) htseq-count with parameters “-m intersection-strict -s no”. The count files were used as input to R package DESeq (Anders and Huber, 2010) for normalization and the differential expression genes were set at a *p* value cutoff of 0.05. The venn diagrams of overlapped differential genes were generated by VENNY-2.1 (http://bioinfogp.cnb.csic.es/tools/venny/index.html). Go term analysis was performed by David (Huang da et al., 2009a, b). WGCNA (1.51) was used for network analysis of co-regulated genes with parameters “Soft thresholding power β=8; Scale-free topology fit index R^2^=0.897). For selected module, Cytoscape (3.5.1) was used for network mapping.

#### Metabolome

The raw MS data (.wiff) were converted to mzML files using ProteoWizard MSConvert (version 3.0.6526) and processed using XCMS (Smith et al., 2006) for feature detection, retention time correction and alignment. Part of XCMS processing parameters were optimized and set as followings: mass accuracy in peak detection = 25 ppm; peak width c = (5, 20); snthresh = 6; bw = 10; minfrac = 0.5. R package CAMERA (Kuhl et al., 2012) was used for peak annotation after XCMS data processing. A customized R script was used to preprocess and extract the MS/MS data for database match with in-house library. The resulting raw data was then preprocessed for data normalization using SVR algorithm (Shen et al., 2016). Pathway enrichment analysis was executed using MetaboAnalyst 3.0 (Xia et al., 2015). In MetaboAnalyst 3.0, metabolic pathways for *Drosophila melanogaster* were selected. The pathway analysis algorithms were hypergeometric test for over representation analysis and relative-betweeness centrality for pathway topology analysis. For normalized intensity calculation, the preprocessed metabolomic data was further scaled (see below).

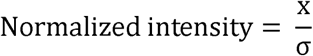

σ is the standard deviation of the population.

Specifically, in each sample group, the pathway intensity was determined by calculating the sum of detected metabolites’ normalized intensities in each metabolic pathway.

## QUANTIFICATION AND STATISTICAL ANALYSIS

All statistical details of experiments are included in figure legends. Sample numbers and experiment replicates are stated in figure legends.

## DATA AND SOFTWARE AVAILABILITY

### Data resources

The raw data files of sequencing experiments have been deposited in the NCBI Gene Expression Omnibus, as well as the normalized read density profiles of ChIP-seq and differential expression results from DESeq of RNA-seq reported in this paper. The accession number is GEO:GSE96654.

